# Neural correlates of phonetic categorization under auditory (phoneme) and visual (grapheme) modalities

**DOI:** 10.1101/2024.07.24.604940

**Authors:** Gavin M. Bidelman, Ashleigh York, Claire Pearson

## Abstract

We tested whether the neural mechanisms of phonetic categorization are specific to speech sounds or generalize to graphemes (i.e., visual letters) of the same phonetic label. Given that linguistic experience shapes categorical processing, and letter-speech sound matching plays a crucial role during early reading acquisition, we hypothesized sound phoneme and visual grapheme tokens representing the same linguistic identity might recruit common neural substrates, despite originating from different sensory modalities. Behavioral and neuroelectric brain responses (ERPs) were acquired as participants categorized stimuli from sound (phoneme) and homologous letter (grapheme) continua each spanning a /da/ - /ga/ gradient. Behaviorally, listeners were faster and showed stronger categorization of phoneme compared to graphemes. At the neural level, multidimensional scaling of the EEG revealed responses self-organized in a categorial fashion such that tokens clustered within their respective modality beginning ∼150-250 ms after stimulus onset. Source-resolved ERPs further revealed modality-specific and overlapping brain regions supporting phonetic categorization. Left inferior frontal gyrus and auditory cortex showed stronger responses for sound category members compared to phonetically ambiguous tokens, whereas early visual cortices paralleled this categorical organization for graphemes. Auditory and visual categorization also recruited common visual association areas in extrastriate cortex but in opposite hemispheres (auditory = left; visual=right). Our findings reveal both auditory and visual sensory cortex supports categorical organization for phonetic labels within their respective modalities. However, a partial overlap in phoneme and grapheme processing among occipital brain areas implies the presence of an isomorphic, domain-general mapping for phonetic categories in dorsal visual system.

## Introduction

The seemingly trivial task of comprehending speech requires the brain to categorize incoming stimulus features into discrete chunks. Like most sensory phenomena, speech poses the problem of invariance in which segments of continuous stimulus features map to discrete categories (Goldstone and Hendrickson, 2010). In speech perception, categorical processing can be inferred from tasks where listeners hear sounds along a morphed phonetic continuum (e.g., /ba/-/pa/) and are asked to label those sounds with a binary response. Typically, listeners perceive the same phoneme until reaching the midpoint of the continuum where their labeling abruptly shifts—the category boundary (Liberman et al., 1967; Pisoni, 1973; Harnad, 1987; Pisoni and Luce, 1987). The categorical division of the speech signal enables interpretation of the continuous acoustic stream as a sequence of phonemes that comprise words and form the basis of subsequent high-order linguistic units. And while originally thought to be unique to speech, several studies have shown that even non-speech stimuli are perceived in a categorial manner including music (Locke and Kellar, 1973; Siegel and Siegel, 1977; Burns and Ward, 1978; Zatorre and Halpern, 1979; Howard et al., 1992; Burns and Campbell, 1994; Klein and Zatorre, 2011; Mankel et al., 2022), colors (Fonteneau and Davidoff, 2007; Franklin et al., 2008), faces (Beale and Keil, 1995), and lines (Foster, 1983; Ferraro and Foster, 1986; Foster and Ferraro, 1989). Thus, both auditory and visual features are supported by a categorical coding scheme in the brain’s perceptual-cognitive system.

Language itself exerts strong influences on audiovisual category representations. For example, relative to English speakers, Chinese speakers show sharper categorization of Mandarin tones (Bidelman and Lee, 2015), and nonmusicians show sharper categorization for native speech sounds relative to unfamiliar musical sounds (Bidelman and Walker, 2017). Linguistic experience also influences categorization of visual information. For example, cultures that distinguish color terms lexically (e.g., shades of blue) show more precise categorization of color spectra (e.g., blue-green) (Winawer et al., 2007). The perception of visual Chinese characters also varies according to ones’ experience with different Chinese orthographic writing systems (Yang and Wang, 2018). Language is therefore tightly coupled with perception and influences the categorical processing of sound and visual information alike.

In this vein, studies have shown that letter-speech sound integration is crucial for reading acquisition (Preston et al., 2016). Interestingly, more experienced readers show sharper perception of phonetic boundaries than poor readers (Werker and Tees, 1987; Mody et al., 1997) and there is robust evidence that the relationship between reading ability and phonological awareness is reciprocal—the ability to isolate phonemes improves with exposure to the alphabet, and reading ability improves with training on phoneme segmentation (for review, see Bentin, 1992). The link between reading and spoken language experience is therefore a critical component of language development. Humans acquire spoken language from birth, master its basic structure by age 3, and do not acquire written language until years later (Miller and Gildea, 1987). Given that it is a less sophisticated skill, early stages of the reading process must transform the visual input to a form that is compatible with the acoustic speech perception system to promote efficient comprehension (Godfrey et al., 1981). Presumably, this involves converting individual graphemes to phoneme-like internal representations, and then mapping sequences of graphemes to encoded representations of syllables and words (Liberman et al., 1967).

Although grapheme and phoneme representations are both critical to language processing, auditory and visual categories might be subject to different degrees of categorical organization in the brain. One study found categorical processing of uppercase letters from a V/X continuum (Yasuhara and Kuklinski, 1978), suggesting linguistic experience with stimulus labels results in letters becoming perceptual wholes. However, other work has shown the perception of letters from a G/Q continuum changes more gradiently as a function of letter ambiguity rather than letter category (Massaro and Hary, 1986). Similarly, another study found that the peak in discrimination performance for an n/h continuum did not correspond with the category boundary (McIntyre and Di Lollo, 1991). These conflicting findings suggest visual graphemes might be perceived more continuously than their sound phoneme counterparts.

While studies of letter categorization have provided conflicting results, other studies have shown categorical processing of visual stimuli that are highly relevant to letter decoding—namely lines. Foster (1983) employed visual stimuli from a curved line continuum in a discrimination task in which participants identified which of four lines was different. The stimuli were employed in an identification task in which participants labeled items as ‘straight’, ‘just curved’, or ‘more than curved’. Peaks in performance from the discrimination task correlated with perceived category boundaries in the identification task (Foster, 1983). Another study of line perception employed items from a curved line continuum in a three alternative forced choice task and found that discrimination performance peaked at the midpoint of the continuum, consistent with category representation (Ferraro and Foster, 1986). Similarly, Foster and Ferraro (1989) examined categorization for visual stimuli from a horizontal/vertical line offset continuum. They found that peaks in performance from a line position discrimination task correlated with category boundaries identified in a labeling task (‘no gap’, ‘gap’, ‘more than just a gap’). These findings suggest the visual features inherent to letter objects (i.e., curved, horizontal, and vertical lines) are indeed perceived in a categorical manner. Letters vary along these and other visual dimensions (e.g., obliqueness) across various fonts and handwriting styles. Although like speech sounds, visual letters represent the basic elements of language and meaning, no studies have directly compared their perceptual categorization. Contrasting letter and speech processing could elucidate whether the neural mechanisms underlying phonological linguistic categories are specific to input sensory modality.

To this end, the aim of the present study was to directly compare speech (phoneme) and letter (grapheme) perception to determine whether the brain employs analogous neural processes across modalities to map continuous features of auditory and visual signals into their discrete linguistic categories. As far as we are aware, no studies have directly compared phoneme (sound) and grapheme (visual) category mapping to identical phonetic units in a cross-modal design. To measure the categorical processing of letters and speech sounds, we recorded multichannel event-related brain potentials (ERPs) as listeners actively categorized stimuli along a “da-ga” phoneme and homologous “da-ga” grapheme continuum. Comparing ERPs to letters and speech sounds allowed us to (i) assess where/when linguistic category representations in each modality emerge in the brain and (ii) distinguish shared vs. segregated neural mechanisms in processing audiovisual analogues that share identical category identity.

## Materials & Methods

### Participants

N=16 young adults (3 male, 13 female; age: *M* = 24.5, *SD* = 12.9 years) participated in the experiment^1^. All exhibited normal hearing sensitivity confirmed via audiometric screening (i.e., < 25 dB HL, octave frequencies 250 - 8000 Hz) and had normal or corrected-to-normal vision. Each participant was strongly right-handed (74.8 ± 27.0% laterality index; Oldfield, 1971), had obtained a collegiate level of education (18.8 ± 2.7 years formal schooling), and was a native speaker of American English. On average, the sample had 3.25 ± 3.3 years of music training. All were paid for their time and gave informed consent in compliance with a protocol approved by the Institutional Review Board at the University of Memphis.

### Auditory (phoneme) and visual (grapheme) stimulus continuum

We used a 5-step, stop-consonant /da/ to /ga/ sound continuum (varying in place of articulation) to assess CP for speech (e.g., Bidelman et al., 2019) (**Fig. 1A**). Each sound token (Tk) was separated by equidistant steps acoustically yet was perceived categorically from /da/ to /ga/. Stimulus morphing was achieved by altering the F2 formant region in a stepwise fashion using STRAIGHT (Kawahara et al., 2008). The original audio material for the /da/ and /ga/ endpoint exemplars were recorded by Nath and Beauchamp (2012). We chose a consonant-vowel (CV) continuum because compared to other speech sounds (e.g., vowels), CVs are perceived more categorically (Pisoni, 1973; Altmann et al., 2014) and carry more salient articulatory gestures and visual cues for perception (Moradi et al., 2017). All tokens were normalized in duration (500 ms), amplitude (75 dB SPL), and bandwidth (50-4000 Hz). The auditory stimuli were delivered binaurally through shielded insert earphones (ER-2; Etymotic Research) controlled by a TDT RP2 signal processor (Tucker Davis Technologies).

**Figure 1:**
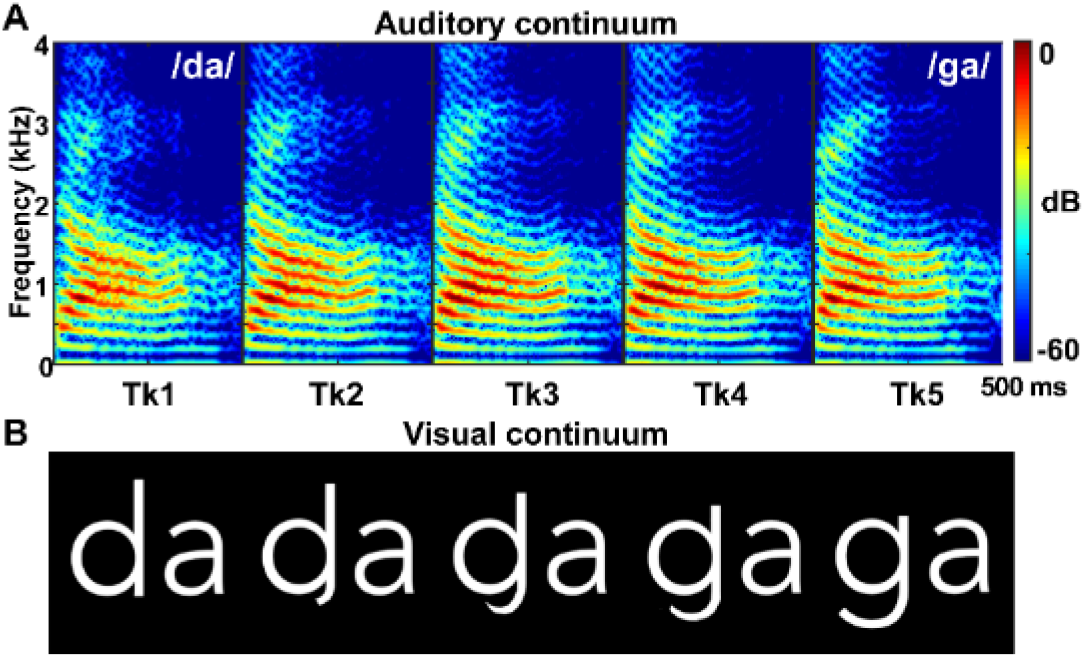
Auditory and visual stimulus continua. **(A)** Phoneme sound continuum spanning 5 equidistant steps between “da” and “ga.” Morphing was achieved by altering the F2 formant region in a stepwise fashion. (**B**) Visual grapheme continuum spanning 5 equidistant steps between “da” and “ga”.

The visual grapheme continuum was created by morphing (5 steps) between the alphanumeric images of “da” and “ga.” Visual morphing was achieved using custom scripts coded in MATLAB (e.g., Jonathan, 2011) (**Fig. 1B**).

During EEG recording, listeners heard 150 trials of each individual token in auditory or visual blocks and labelled the stimulus with a binary response (“da” or “ga”) as quickly and accurately as possible on the computer keyboard. Following, the interstimulus interval (ISI) was jittered randomly between 800 and 1000 ms (20 ms steps, uniform distribution) to avoid rhythmic entrainment of the EEG and anticipating subsequent stimuli. Block order for modality was randomized within and between participants. Visual stimuli were presented at the center of the computer screen (Samsung SyncMaster S24B350HL; nominal 75 Hz refresh rate) on a black background at a distance of ∼1 m.

### EEG recordings

EEGs were recorded from 64 sintered Ag/AgCl electrodes at standard 10-10 scalp locations (Oostenveld and Praamstra, 2001). Continuous data were digitized at 500 Hz (SynAmps RT amplifiers; Compumedics Neuroscan) using an online passband of DC-200 Hz. Electrodes placed on the outer canthi of the eyes and the superior and inferior orbit monitored ocular movements. Contact impedances were maintained < 10 kΩ. During acquisition, electrodes were referenced to an additional sensor placed ∼ 1 cm posterior to Cz. Data were re-referenced offline to the common average for analysis. Pre-processing was performed in BESA® Research (v7.1) (BESA, GmbH). Ocular artifacts (saccades and blinks) were corrected in the continuous EEG using principal component analysis (PCA) (Picton et al., 2000). Remaining trials exceeding ±150 μV were further discarded. Cleaned EEGs were then filtered (1-20 Hz), epoched (−200-800 ms), baselined to the pre-stimulus interval, and ensemble averaged resulting in 10 ERP waveforms per participant (5 tokens*2 modalities).

### Behavioral data analysis

Identification scores were fit with a sigmoid function *P* = 1/[1+*e*^-β*1*(*x* - β*0*)^], where *P* is the proportion of trials identified as a given phoneme, *x* is the step number along the stimulus continuum, and β_*0*_ and β_*1*_ the location and slope of the logistic fit estimated using nonlinear least-squares regression. Comparing parameters between speech contexts revealed possible differences in the “steepness” (i.e., rate of change) of participant’s category labeling in the auditory and visual modality. Steeper functions represent stronger binary categorization. Behavioral labeling speeds (i.e., reaction times [RTs]) were computed as listeners’ trimmed median response latency across trials for a given condition. RTs outside 250-2500 ms were deemed outliers (e.g., fast guesses, lapses of attention) and were excluded from the analysis (Bidelman et al., 2013; Bidelman and Walker, 2017).

### EEG data analysis

#### ERP peak analysis

We measured the amplitude and latency of the auditory evoked potential (AEP) P2 deflection between 175-250 ms. We focus on the auditory P2 as we have previously shown this defection indexes auditory object and speech identification (Bidelman et al., 2013; Bidelman and Walker, 2019; Bidelman et al., 2020; Bidelman et al., 2021) and tracks with perceptual learning during auditory categorization tasks (Mankel et al., 2022; MacLean et al., 2024). Similarly, we measured the peak positivity from visual evoked potentials (VEPs) within the 375-450 ms time window. These analysis windows were guided by visual inspection of the grand averaged data which showed peak activation in this timeframe (see Fig. 3).

#### Topographic ANOVA (TANOVA)

To provide a more comprehensive analysis of where effects emerged over the entirety of time and space, we used a topographic ANOVA (TANOVA) to identify the spatiotemporal points where the ERPs were sensitive to our stimulus manipulations (i.e., token and modality) (for details, see Murray et al., 2008; Koenig and Melie-Garcia, 2010; Bidelman and Yellamsetty, 2017). TANOVAs were implemented in the MATLAB package Ragu (Koenig et al., 2011; Habermann et al., 2018).

The TANOVA used a randomization procedure (N=500 resamples) that tested the distribution of the ERP’s topography in the measured data against a surrogate distribution, derived by exchanging all conditions and electrodes in the data. The percentage of shuffled cases where the effect size obtained after randomization was equal to or larger than the measured effect size obtained in the observed data provided an estimate of the probability of the null hypothesis. This analysis yielded running *p*-values across the epoch that identified the time samples at which the ERPs were significantly modulated by main (modality, token) and interaction effects (modality x token). To be considered a reliable effect (and prevent Type I error inflation), the procedure required a duration threshold whereby ≥46 ms of contiguous samples had to survive a *p*<0.05 criterion to be considered a significant time window.

From the running TANOVA, we identified time segments where the ERPs showed modulations with both stimulus factors (i.e., modality x token interaction). One such window (centered at ∼200 ms) survived correction for multiple comparisons (see *, Fig. 5A). Within this window, differences in factor levels were visualized by computing ERP difference maps (averaged across the window’s duration) between the scalp topographies for each condition. Because the number of case dimensions is large (e.g., 64 sensors x 2 modalities x 5 tokens x 15 subjects), the visualization was reduced to a two-dimensional space using a multidimensional scaling (MDS) approach (Koenig et al., 2011). Similarities between the mean scalp topographies of the different conditions were assessed using the covariance between maps. The two-dimensional space that optimally represented the covariance matrix was represented in the first two eigenvectors. MDS visualization was then achieved by projecting the mean scalp map for a given condition/factor level onto the two eigenvectors, yielding a set of two-dimensional coordinates of each mean different scalp field map that was displayed as a scatterplot (see Fig. 5B). Points closer in the MDS neural space indicate a high degree of similarity between the scalp topographies whereas farther points reflect more dissimilar neural responses (Bidelman et al., 2013).

#### Source imaging analysis

To estimate the underlying sources contributing to categorial processing, we used Standardized Low Resolution Electromagnetic Tomography (sLORETA) (Pascual-Marqui, 2002) to estimate the neuronal current density underlying the scalp ERPs. This distributed inverse method uses a standardized, unweighted minimum norm. sLORETA models the inverse solution as a large collection of elementary dipoles distributed over nodes on a mesh of the cortical volume. The algorithm estimates the total variance of the scalp data and applies a smoothness constraint to ensure current changes minimally between adjacent brain regions (Picton et al., 1999; Michel et al., 2004). sLORETA source images were computed in the 154-256 ms time window, where modality x token interaction effects were prominent in the TANOVA of the scalp data (see Fig. 5).

## Results

Behavioral identification functions are shown for phoneme and grapheme continua in **Figure 2**. Listeners showed stair-stepped labeling in both stimulus modalities confirming a sharp flip in their category percept from “da” to “ga” at the midpoint of each continuum (**Fig 2. A, B**). A 1-way ANOVA revealed identification slopes were steeper for auditory compared to visual stimuli [*F*_*1,14*_ = 90.11, *p* < 0.0001; = 0.87] indicating stronger categorial hearing of sounds than homologous visual tokens.

**Figure 2:**
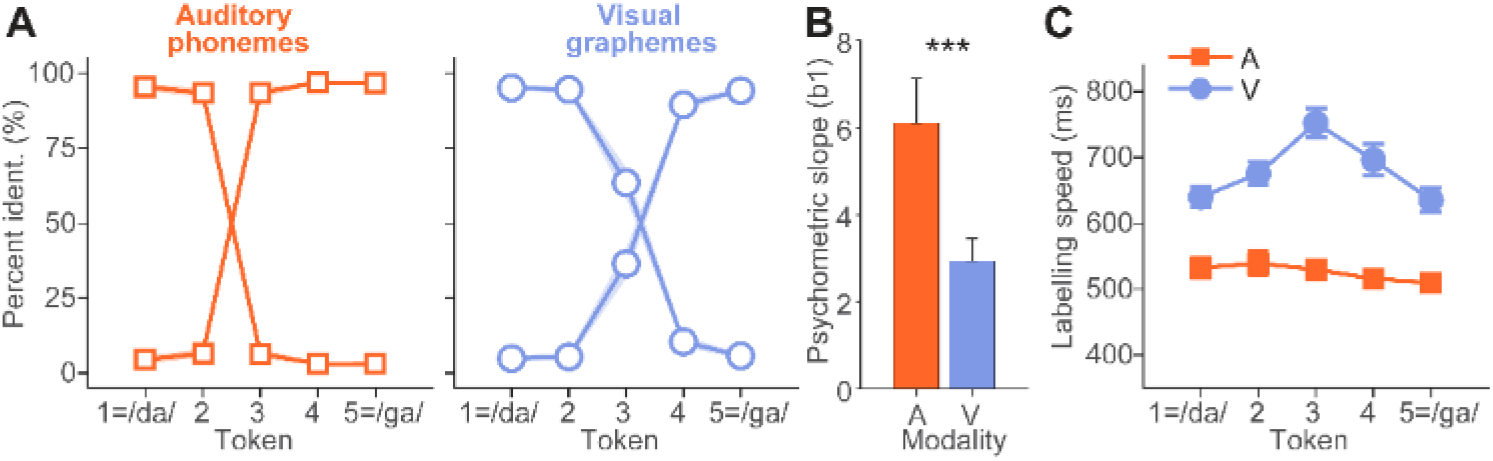
Behavioral categorization of auditory phonemes and visual graphemes of a /da/-/ga/ continuum. **(A)** Identification functions. A and V token labeling shows a stair-stepped identification function content with categorical hearing; listeners perception abruptly flips at the midpoint of the continua (i.e., categorical boundary). (**B**) Slopes of the psychometric functions for auditory and visual categorization. Participants showed sharper (i.e., more categorical) perception of A vs. V tokens. (**C**) Reaction time (RT) speeds for token labelling. Responses are faster for A than V stimuli overall. Visual stimuli also showed a slowing near the midpoint vs. endpoint tokens, consistent with category ambiguity near the midpoint of the continuum (Pisoni and Tash, 1974). Errorbars = ±1.s.e.m., \*\*\**p* < 0.0001

**Figure 3:**
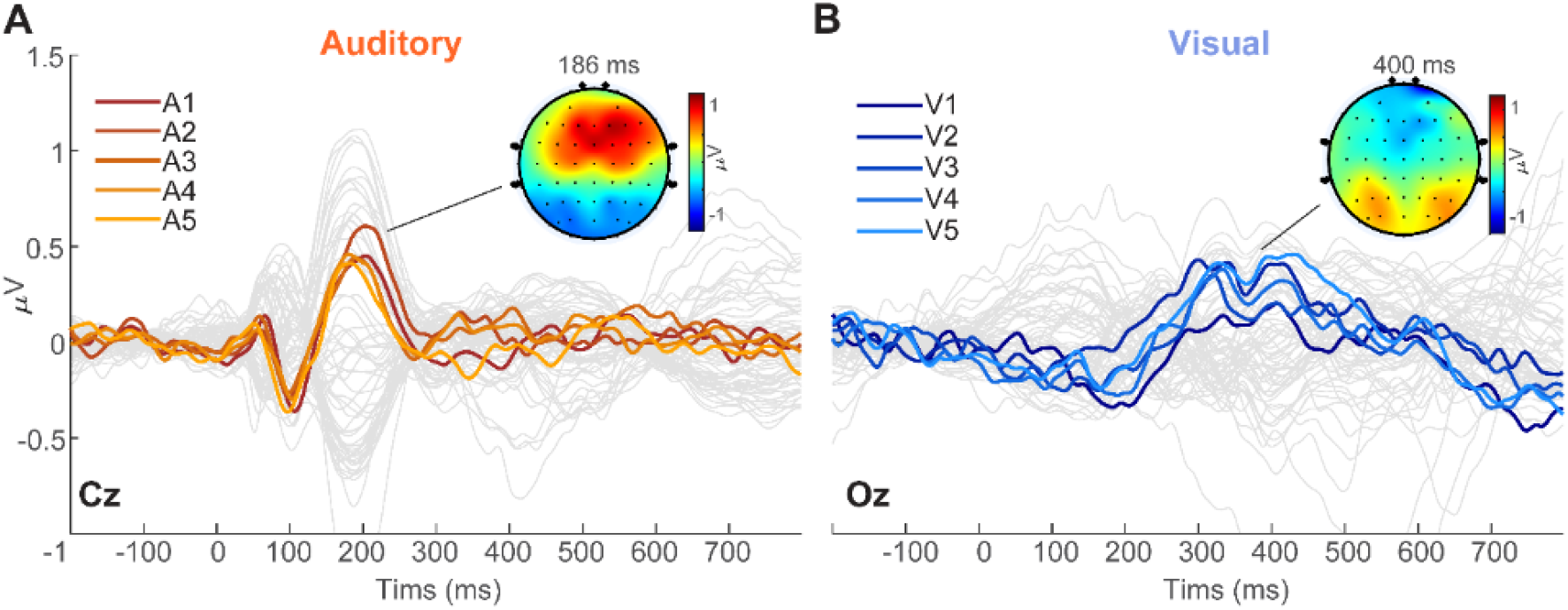
Grand average cortical brain responses to phoneme and grapheme stimuli along a /da/-/ga/ continuum. **(A)** AEPs recorded at the vertex (Cz electrode). Auditory responses reveal a canonical P1-N1-P2 response with frontocentral topography. Note the token-related modulation in the time window of the P2 (∼200 ms) (**B**) VEPs recorded at posterior scalp (Oz electrode). VEPs were weaker than their AEP counterparts overall and showed stimulus-related modulations later in time (300-400 ms). Topographic maps are pooled across tokens for each modality.

RTs were highly sensitive to both stimulus manipulations. Decision speeds showed a strong effects of modality [*F*_*1,126*_ =424.04, *p*<0.0001; = 0.2077 and token [*F*_*4,126*_ =9.53, *p*<0.0001; = 0.23], but more critically, a modality x token interaction [*F*_*4,126*_ =7.74, *p*<0.0001; = 0.20] (**Fig. 1C**). The interaction was attributed to a slowing of RT speeds near the midpoint of the continuum where category membership becomes perceptually ambiguous (Pisoni and Tash, 1974; Bidelman and Walker, 2017; Bidelman and Carter, 2023). This inverted V-shape pattern in RTs was observed only for the V (contrast Tk3 vs. mean of others *t*_*126*_=6.84, *p* < 0.0001) but not the A continuum (*t*_*126*_=0.413, *p* =0.68). Taken together, the overall sharper categorical pattern and faster overall RTs for A vs. V tokens suggests stronger categorization for auditory phoneme compared to visual grapheme stimuli.

Grand average auditory- (AEPs) and visual-(VEP) evoked potentials elicited by phoneme and graphemes, respectively, are shown in **Figure 3**. AEPs revealed a canonical P1-N1-P2 response with frontocentral topography that flipped in polarity at the mastoids— consistent with neural generators in the supratemporal plane (Picton et al., 1999). Token-related modulations were observed in the time window of the P2 (∼200 ms), consistent with the notion this wave reflects auditory object and speech categorization (Bidelman et al., 2013; Bidelman and Walker, 2019; Bidelman et al., 2020; Bidelman et al., 2021; Mankel et al., 2022; MacLean et al., 2024). In contrast, VEPs were maximal at the posterior of the scalp, consistent with generators in the visual cortices surrounding the calcarine fissure (Ducati et al., 1988). VEPs were generally weaker than their AEP counterparts overall and showed stimulus-related modulations later in time (300-400 ms).

A mixed-model ANOVA conducted on ERP peak amplitudes showed no effects of modality or token alone [all *ps >* 0.23] (**Fig. 4A**). In stark contrast, ERP latencies showed a main effect of stimulus modality that paralleled the behavioral RTs; VEPs were later than their AEP counterparts across the board [*F*_*1,126*_ = 3082.89, *p* < 0.0001; = 0.96]. However, this effect was expected given the difference in analysis window and peak deflection across AEP and VEP waveforms. More critically, ERP latencies showed a modality x token interaction [*F*_*4,126*_ = 2.82, *p* = 0.027; = 0.08] (**Fig. 4B**). The interaction suggests that the degree of categorical coding across tokens varied by stimulus modality. Supporting this notion, a repeated measures correlation (rmCorr) (Bakdash and Marusich, 2017) across tokens and modalities showed ERP latency was positively correlated with behavioral RTs (*r*_*rm*_ = 0.79, *p* < 0.0001). Faster neural latencies were associated with faster categorization speeds, which was more prominent in the auditory than visual modality.

**Figure 4:**
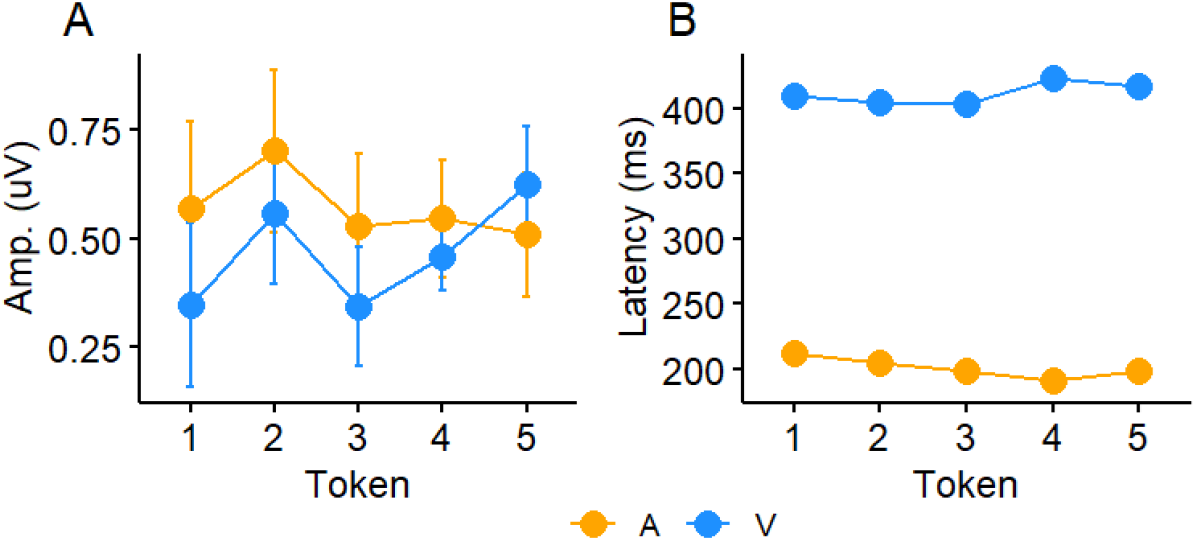
ERP (A) amplitude and (B) latency as a function of stimulus modality and CV token. Errorbars = ±1.s.e.m.

**Figure 5:**
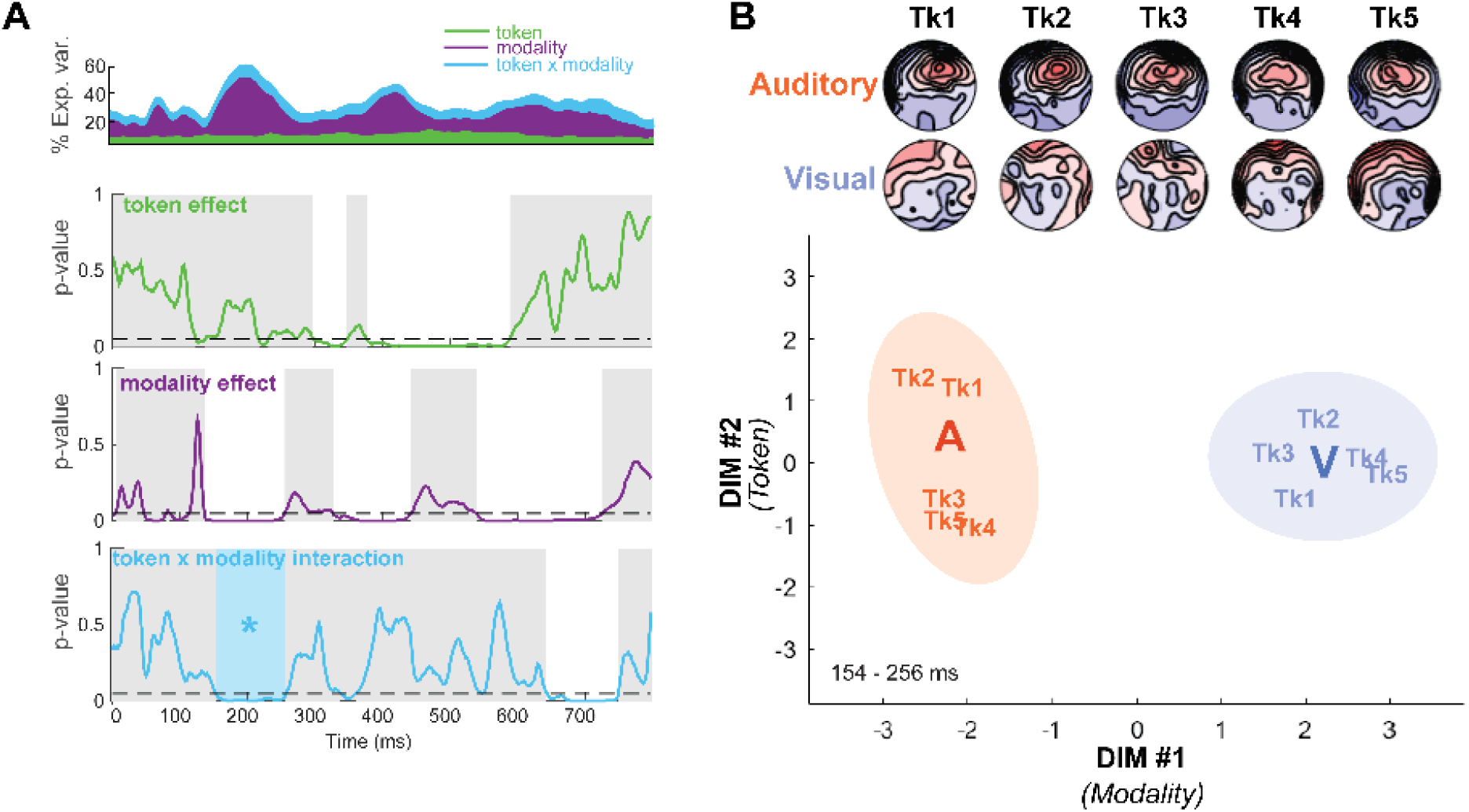
Topographic ANOVA (TANOVA) revealing the time course at which the brain distinguishes phoneme and grapheme tokens. (**A**) *Top*, running time course of the %-explained variance of the TANOVA over all electrodes and time for the main and interaction effects. *Bottom 3 rows*, time course for the main effect of token, modality, and token x modality interaction. Each trace represents the running *p*-value for the effect, computed via permutation resampling (N=500 shuffles). Dotted lines mark the *p*=0.05 significance level. \**p* < 0.05 (corrected), shaded regions= *n*.*s*. (**B**) MDS visualization of the token x modality interaction (i.e., see* panel A). *Top*, scalp topographies per token and modality. *Bottom*, MDS visualization of the differences between neural responses to phoneme and grapheme stimuli. Note the clear separation of responses to A and V tokens along dimension #1 (*modality*) and clustering of within-category tokens (e.g., adjacent Tk1/Tk2 and far from Tk4/Tk5) within each modality. Category clustering of phonemes/graphemes appears more prominent in the auditory than visual modality.

TANOVAs (Murray et al., 2008; Koenig et al., 2011) conducted on the ERP topographies confirmed significant modulations in evoked activity with changes in both stimulus modality and token, as well as segments sensitive to both acoustic factors (i.e., modality x token interaction) (**Fig. 5A**). By itself, token modulated activity across the middle portion of the response time course beginning at ∼300 ms post stimulus onset. The main effect of modality was more pervasive, with significant modulations beginning as earlier as ∼150 ms. The significant token x modality interaction was circumscribed to an early (∼154-256 ms) and late (∼700 ms) time window after stimulus onset. In terms of the processes that unfold during the chronometry of perceptual identification, these segments have been described as encoding vs. decision periods of the categorization process (Mahmud et al., 2021). As such, subset analyses focused on the early sensory-encoding window.

Scalp topographies and MDS visualization of the AEP and VEP responses to phoneme and grapheme stimuli are shown in **Fig. 5B**. Responses clustered into two distinct “clouds” in the MDS based on stimulus modality (A vs. V: DIM #1). Similarly, individual tokens showed differentiation along the orthogonal DIM #2. Consistent with prior studies examining the neural differentiation of phonetic categories in AEPs (Chang et al., 2010; Bidelman et al., 2013; Bidelman and Lee, 2015), within-category tokens (e.g., Tk1/2) tended to cluster in closer proximity to one another in the MDS space but far from their across-category counterparts (e.g., Tk 4/5). More critically, tokens grouped within their respective modality, as indicated by the token x modality interaction observed in the TANOVA. However, category clustering of phonemes/graphemes appeared to be more prominent in the auditory than visual modality as indicated by a tighter convergence of within-category stimuli and farther separation across categories, respectively.

To resolve the neuronal sources underlying these modality-specific responses during categorization, we used sLORETA imaging (Pascual-Marqui, 2002) to visualize the current densities on the cortical surface. Statistical maps were generated contrasting the degree of categoricity in the neural AEP and VEP responses. To this end, we first computed difference waves between the two prototypical (i.e., mean Tk1/5) and phonetically ambiguous (i.e., Tk 3) tokens (separately for A and V responses). These difference waves index the degree of categorial coding in the ERPs (Liebenthal et al., 2010; Bidelman and Walker, 2017; Bidelman and Walker, 2019). We then calculated *t*-statistic maps contrasting this categorical coding index between modalities [i.e., (A_Tk1/5_ – A_Tk3_) – (V_Tk1/5_ – V_Tk3_)]. Maps were computed in the 154-256 ms time window, where responses showed a maximal token x modality interaction (i.e., Fig. 5). The resulting statistical maps compared the degree of categorical coding in the auditory vs. visual modality across the entire brain volume.

Source activations for phoneme vs. grapheme categorization are shown in **Fig. 6**. Category coding for speech sound phonemes was stronger than visual graphemes in bilateral auditory cortex (AC), left inferior frontal gyrus (IFG), and middle frontal gyrus (MFG). Surprisingly, auditory categorical coding more strongly recruited portions of left occipital cortex including extrastriate visual association areas (BA 18/19). In contrast, visual categories recruited a network involving nodes in bilateral precentral gyrus (PCG), cuneus (CUN), and the right dorsal stream of the visual pathway including lateral occipital cortex and visual association areas (BA 18/19).

**Figure 6:**
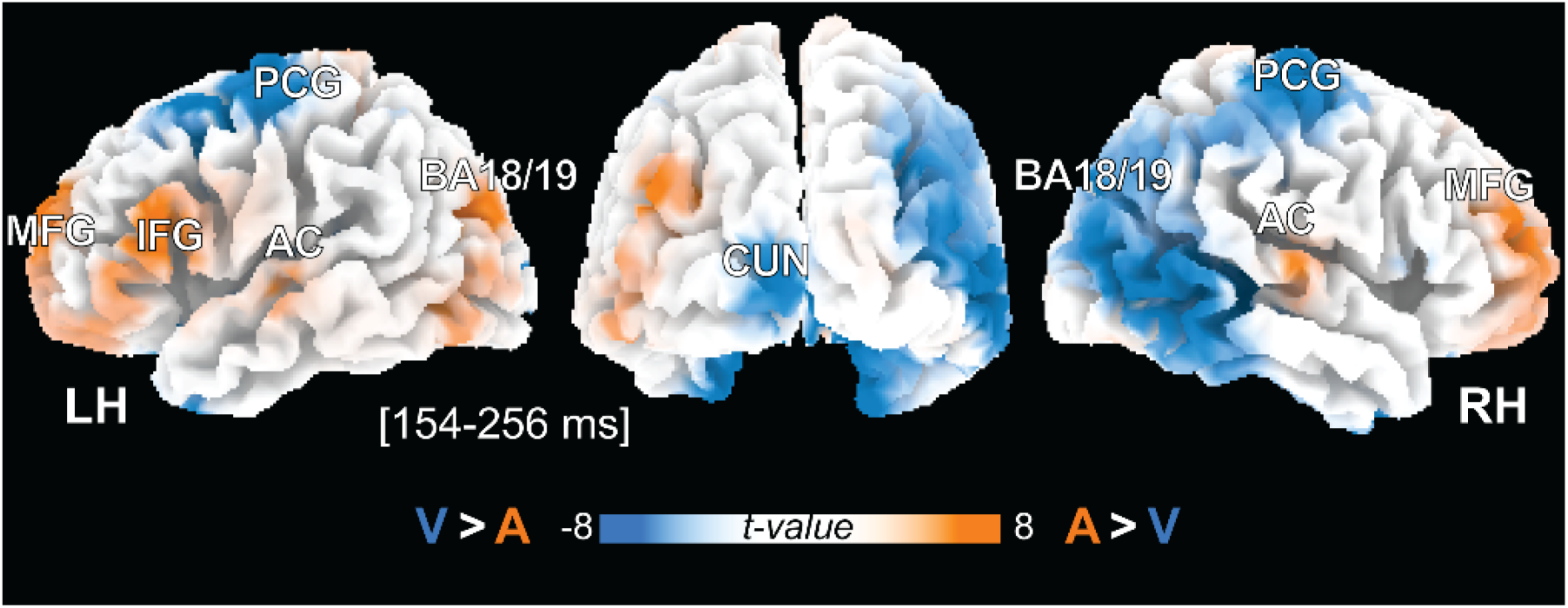
Source responses reveal differential activation of auditory, visual, and linguistic cortex during audiovisual categorization that depend on stimulus modality. sLORETA statistical maps between A and V responses (*t*-stat, p < 0.05 masked, corrected) projected onto the Collins brain template (Collins et al., 1998). The contrast reflects the difference in degree of categorical coding between the auditory and visual modalities [i.e., (A_Tk1/5_ – A_Tk3_) – (V_Tk1/5_ – V_Tk3_)]. Maps are shown in the 154-256 ms time window during the token x modality interaction (see Fig. 5A). Hot colors denote preferential coding of auditory categories; cool colors, preferential coding of visual categories. AC, auditory cortex; BA, Brodmann area; CUN, cuneus; MFG, middle frontal gyrus; IFG, inferior frontal gyrus; PCG, precentral gyrus; LH/RH, left/right hemisphere.

## Discussion

A key aspect of language comprehension is the ability to categorize both acoustic and written inputs to form discrete linguistic units. Understanding how different sensory modalities perform this mapping between stimulus features and abstract linguistic space is important for understanding human speech comprehension. By measuring behavioral and neuroelectric brain responses (ERPs) during a sound (phoneme) and letter (grapheme) /da/ - /ga/ continuum, our data reveal (i) both modality-specific and overlapping brain regions support cross-modal categorization and (ii) stronger categorization for speech phonemes than their homologous orthographic counterparts. Collectively, our results imply that acoustic information enjoys a privileged role in the perceptual-cognitive operation of categorization.

### Categorization is more salient for auditory than visual linguistic stimuli

Behaviorally, we found both phonemes and graphemes elicited the typical, stair-stepped identification functions characteristic of categorical hearing (Pisoni, 1973). Whether or not letters are perceived categorically has been somewhat equivocal in the literature (Yasuhara and Kuklinski, 1978; Massaro and Hary, 1986). Our data extend prior studies on visual objects (e.g., lines, colors, and faces; Foster, 1983; Ferraro and Foster, 1986; Foster and Ferraro, 1989; Beale and Keil, 1995; Fonteneau and Davidoff, 2007; Franklin et al., 2008) by confirming a similar category mapping for orthographic CV letters. However, we show here that both auditory and visual CV homologues are perceived in a categorical manner but to differing degrees. Listeners were faster and showed stronger, more discrete labeling of phoneme compared to grapheme CV tokens. Moreover, labeling was slower overall for visual than auditory stimuli and graphemes showed an additional slowing near the continuum’s midpoint which was not observed for phoneme tokens. Our identification and RT data suggest that visual graphemes were perceived less categorically (i.e., more continuously) and/or were more perceptually ambiguous than their auditory counterparts (Pisoni and Tash, 1974; Bidelman and Walker, 2017; Carter et al., 2022; Bidelman and Carter, 2023; Rizzi and Bidelman, 2024). Taken together, the overall sharper categorical pattern and faster overall RTs for auditory vs. visual tokens suggests stronger categorization for auditory phonemes compared to visual grapheme stimuli.

Paralleling the behavioral data, we found faster neural timing corresponded with faster perceptual categorization speeds. ERP responses also peaked ∼200 ms earlier for auditory compared to visual stimuli. Moreover, MDS scaling of the EEG revealed responses self-organized in a categorial fashion such that tokens clustered within their respective modality beginning ∼150-250 ms after stimulus onset. These data are consistent with prior studies examining the neural differentiation of phonetic categories in the AEPs (Chang et al., 2010; Bidelman et al., 2013; Bidelman and Lee, 2015), where within-category tokens (e.g., Tk1/2) tend to cluster in closer proximity to one another but far from their across-category counterparts (e.g., Tk 4/5). Category clustering was also more prominent in the auditory than visual modality as indicated by a tighter convergence of within-category stimuli and farther separation across categories for phonemes. Collectively, our behavioral and neuroimaging data suggest the neural differentiation and subsequent grouping of tokens into categorical representations is more binary and robust in the auditory vs. visual modality for stimuli otherwise matched in phonetic identity.

Although our study only assessed uni-sensory audio/visual responses, our results extend a large body of work on multisensory integration in categorization. Integrating multiple cues is necessary in face-to-face communication in which visual articulatory information from a talker’s face provides a critical complement to what was said. In audiovisual contexts, dynamic speech features in auditory and visual channels reflect discrete representations of phonetic-linguistic units (phonemes) and corresponding representations of mouth shapes (visemes) (Peelle and Sommers, 2015) that can interact to systematically influence the perceptual identity of speech objects themselves (Massaro and Cohen, 1983; van Wassenhove et al., 2005; Bidelman et al., 2019). Electrophysiological studies have also shown that visual stimuli modulate auditory cortical responses in the auditory cortical fields (Kayser et al., 2008) and visual cues can increase the precision of category representations leading to a sharper perceptual division of the speech signal that aids its perception (Bidelman et al., 2019). The activation of primary auditory cortex during lip reading further implies visual cues might influence perception even before speech sounds are categorized into their phonetic constituents (Calvert et al., 1997; Bernstein and Liebenthal, 2014).

Cross-modal interactions within sensory brain regions have also been observed in human neuromagnetic brain responses to auditory and visual stimuli (Raij et al., 2010). These studies reveal that while cross-sensory (auditory→visual) activity generally manifests later (∼10-20 ms) than sensory-specific (auditory→auditory) activations, there is a stark asymmetry in the arrival of information between Heschl’s gyrus and the Calcarine fissure. Auditory information is combined in visual cortex roughly 45 ms faster than the reverse direction of travel (i.e., visual→auditory) (Raij et al., 2010) and auditory cues can bias and “override” normal perception of visual objects (Bidelman and Myers, 2020). Such dominance of auditory compared to visual information in multisensory studies might explain the larger and more extensive categorical organization we find for speech-sound phonemes compared to grapheme equivalents.

### Phoneme and grapheme categorization recruit shared and segregated brain networks

Source reconstruction of the ERPs revealed distributed, but partially overlapping neural networks supporting phoneme vs. grapheme categorization. These findings are broadly consistent with previous functional connectivity studies that have identified a sparse but distributed network supporting phonetic categorization including areas of left linguistic (IFG), visual (cuneus/precuneus), and motor cortex (central gyrus) (Al-Fahad et al., 2020; Mahmud et al., 2021). However, direct comparisons between continuum revealed category coding for speech sound phonemes was stronger than visual graphemes in bilateral auditory cortices, left IFG, and middle frontal gyrus (MFG). Engagement of auditory cortex in processing sound categories is consistent with prior work showing early auditory cortical areas typically associated with sensory-stimulus coding are highly sensitive to the category structure in speech (Chang et al., 2010; Bidelman and Lee, 2015; Bidelman and Walker, 2019; Mankel et al., 2020; Carter and Bidelman, 2021; Rizzi and Bidelman, 2024). Similarly, left IFG has been implicated in phoneme category selectivity (Myers et al., 2009; Alho et al., 2016; Bidelman and Walker, 2019) and resolving ambiguity in the speech signal—as in cases of additive noise or lexical uncertainty (Luthra et al., 2019; Carter and Bidelman, 2021) (but see Hickok et al., 2011). We have also shown left MFG is recruited when listeners experience perceptual shifts in their hearing of speech categories dependent on top-down factors such as stimulus context or lexical biasing (Bidelman et al., 2021; Carter et al., 2022). Left IFG might also be involved in articulatory rehearsal during phonetic perception (Zatorre et al., 1992). Broadly speaking, the engagement of MFG and IFG in our auditory tasks is consistent with the notion that sound categorization recruits post-perceptual processing of the frontal lobes (Binder et al., 2004; Myers et al., 2009; Carter et al., 2022). However, we note these are relatively fast processes, engaging reciprocal auditory-frontal pathways by ∼250 ms after sound enters the ear (see also Bidelman et al., 2021; Mahmud et al., 2021).

In contrast, we found visual categories recruited a network involving nodes in bilateral precentral gyrus (PCG), cuneus (CUN), and the right dorsal stream of the visual pathway including lateral occipital cortex and visual association areas (BA 18/19). These latter regions form the brain’s canonical reading and visual word form areas (Selpien et al., 2015). Stronger engagement of auditory vs. visual cortex for phonemes and graphemes, respectively, is perhaps expected and implies a unitary division for processing modality-specific sensory information. Indeed, as with primary auditory cortex (Chang et al., 2010; Bidelman and Lee, 2015), category-specific information can be read out from early visual cortex (Vetter et al., 2014). Auditory activations have also been shown to predicted phonological processing and rapid automatized naming whereas precuneus activations have been shown to predict reading and writing skills (Xu et al., 2018), respectively. Precentral engagement during visual grapheme tokens is also consistent with prior literature demonstrating activation of primary motor cortex and supplementary motor areas during the perception of handwritten letters (Longcamp et al., 2011).

Surprisingly, we found auditory categories more strongly recruited portions of occipital cortex including extrastriate visual association areas (BA 18/19). These findings are interesting because they suggest nonretinal information is not only coded in the activity patterns of early visual cortex but that sounds might be processed in visual system in a form of speech imagery (Vetter et al., 2014). Indeed, some have argued that the left dominance for language even originates in extrastriate cortex (Selpien et al., 2015). However, we also found these effects depended on input modality and hemisphere. Whereas sound phonemes more strongly recruited *left* BA 18/19, letter graphemes recruited its homologue in *right* hemisphere. This implies a hemispheric asymmetry in the division of labor when processing acoustic vs. visual categories with the same phonetic identity. While the basis of this asymmetry is not fully clear, it interesting to note the lateralization of ventral occipital responses varies with reading expertise (Seghier and Price, 2011). Neural activity is left lateralized for words in skilled readers but right lateralized in novice readers who have not yet learned to link print to sound (Maurer et al., 2006). Conceivably, lateralization might also vary according to individual differences in the related skill of letter-speech sound integration. Under this notion, the stark occipital lateralization we find for auditory (left) vs. visual (right) categorization may result from individual differences in orthographic decoding or auditory–visual matching. At the very least, our results suggest that extrastriate brain areas might perform an isomorphic, domain-general mapping for phonological categories in dorsal visual system. Such a computational hub would allow the brain to map linguistically-relevant sounds and visual linguistic objects alike into a common lexical (rather than purely auditory or visual) representation. Future studies employing audiovisual speech could test this possibility (e.g., Massaro and Cohen, 1983; Bidelman et al., 2019).

## Acknowledgments

Work supported by the National Institutes of Health (NIH/NIDCD R01DC016267). The authors thank Gwyneth Lewis for assistance in data collection.

## Footnotes

1 EEG was not recorded from one participant due to technical error resulting in a final sample size of n=15 for the neural data (behavioral data were unaffected). The current sample was identical to the participants reported in Bidelman et al. (2021).

